# The Transcription Factor Binding Landscape of Mouse Development

**DOI:** 10.1101/2024.08.23.609315

**Authors:** Anna Nordin, Gianluca Zambanini, Mattias Jonasson, Tamina Weiss, Yorick van de Grift, Pierfrancesco Pagella, Claudio Cantù

## Abstract

Gene regulators physically associate to the genome, in a combinatorial fashion, to drive tissue-specific gene expression programs. Uncovering the genome-wide activity of all gene regulators across tissues is therefore needed to understand how the genome is regulated during development. Here, we take a first step forward towards achieving this goal. Using CUT&RUN, we systematically measured the genome-wide binding profiles of key transcription factors and cofactors that mediate the activity of ontogenetically relevant signaling pathways in select mouse tissues at two developmental stages. Computation of the numerous genome-wide binding datasets unveiled a large degree of tissue and time-specific activity for each gene regulator, and several factor-specific idiosyncrasies. Moreover, we identified “popular” regulatory regions that are bound by a multitude of pathway regulators. Popular regions tend to be more evolutionarily conserved, implying their essentiality. Consistently, they lie in the proximity of genes whose dysregulation causes early embryonic lethality in the mouse. Moreover, the human homologs of these regions are also bound by many gene regulators and are highly conserved in human populations, indicating that they retain functional relevance for human development. This work constitutes a decisive step towards understanding how the genome is simultaneously read and used by gene regulators in a holistic fashion and unveils multiple genomic mechanisms that drive embryonic development.

## Main

Transcriptional regulation during embryonic development relies on the tissue-specific deployment of combinatorial systems of DNA-binding transcription factors. Transcription factors physically associate with regulatory regions along the genome, recruiting non-DNA-binding co-factors and chromatin remodelers. Their combined action generates chromatin domains that are permissive for the activity of RNA Polymerase II, leading to transcription (Klemm et al., 2019). Characterizing the genome-wide binding profile of all the different classes of proteins that are involved in this process– that from here on we globally refer to as gene regulators (GRs) – is therefore a necessary step for understanding how the genome is regulated in each cell during development (Merika and Thanos, 2001). As technologies that map the genome-wide physical behavior of these proteins are not trivial to employ, classic experimental approaches typically aim at targeting one or a limited number of such players. Here we propose that, to truly understand how the genome is differentially regulated across cell types during development, one should measure the activity of *all* gene regulators at different developmental time-points. Vertebrates’ genomes, however, encode for ca. 1600 transcription factors and a comparable number of other regulators (Ng et al., 2021). The number and combinatorial activity of GRs render their comprehensive study difficult experimentally. State-of-the-art technologies now allow us to approach this endeavor.

Among the technologies that have been developed to profile the DNA-binding behavior of GRs, Cleavage Under Targets and Release Using Nuclease (CUT&RUN, hereafter C&R) (Skene and Henikoff, 2017; Skene et al., 2018) and CUT&Tagmentation (CUT&Tag; hereafter C&T) are emerging as methods of choice. C&R, in contrast to ChIP-seq, is conducted under native conditions without cross-linking agents – a fact that drastically reduces generation of experimental artifacts (Landt et al., 2012). Moreover, C&R and C&T permit scaling down in cell numbers, with some applications even allowing to yield results from single cells (Bartosovic et al., 2021). While both C&R and C&T reliably work for the detection of the abundant and stable post-translational modifications of histones, they sometimes “suffer” when targeting less stable chromatin regulators such as lowly-expressed transcription factors and non-DNA-binding GRs (Meers et al., 2019). This drawback led us to develop C&R-Low Volume-Urea (LoV-U), that robustly detects all types of GRs, and facilitates intermediate-scale parallel profiling thanks to low sample volumes (Zambanini et al., 2022).

Here, by using C&R-LoV-U (generally referred to as C&R), we initiated the systematic mapping of GRs during mouse organogenesis. We produced a dataset of >100 C&R tracks of GRs acting downstream of developmental signaling pathways in four different mouse tissues (forelimbs (FL), hindlimbs (HL), branchial arches (BA) and liver (L)), at two stages of organogenesis (10.5 and 11.5 days post coitum, dpc). While we realize that this is only the beginning of a larger enterprise, this core dataset allowed us to observe key regulatory mechanisms of development. Our findings include measuring the extent to which each GRs has shared or tissue-specific activity, how GRs deployment changes over time, and the convergence – or crosstalk – of developmental signaling pathways on the genome. Publication of this work is also meant to represent a manifesto testifying for the need for larger scale experimental endeavors. Our hope is that other interested researchers in the field of genome regulation during development will assist us in gathering further resources and experimental power to profile and characterize all GRs.

## Results

### Profiling the genome-wide binding of transcriptional regulators in mouse development

A bird-eye-view is needed to understand how GRs coordinate the complexity of gene expression during development. To initiate the simultaneous profiling of the genome-wide binding behavior of GRs in developing mouse tissues, we decided to perform C&R on tissues dissected from embryonic 11.5 days post coitum (dpc), when organogenesis is established and ongoing (Cao et al., 2019). We selected four tissues for analysis: forelimb buds (FL), hindlimb buds (HL), liver (L), a prominent hematopoietic site at this stage (Jagannathan-Bogdan and Zon, 2013), and branchial arches (BA), through which neural crest cells migrate to contribute to facial and trunk processes (Hari et al., 2012) (Figure 1A).

**Figure 1.**
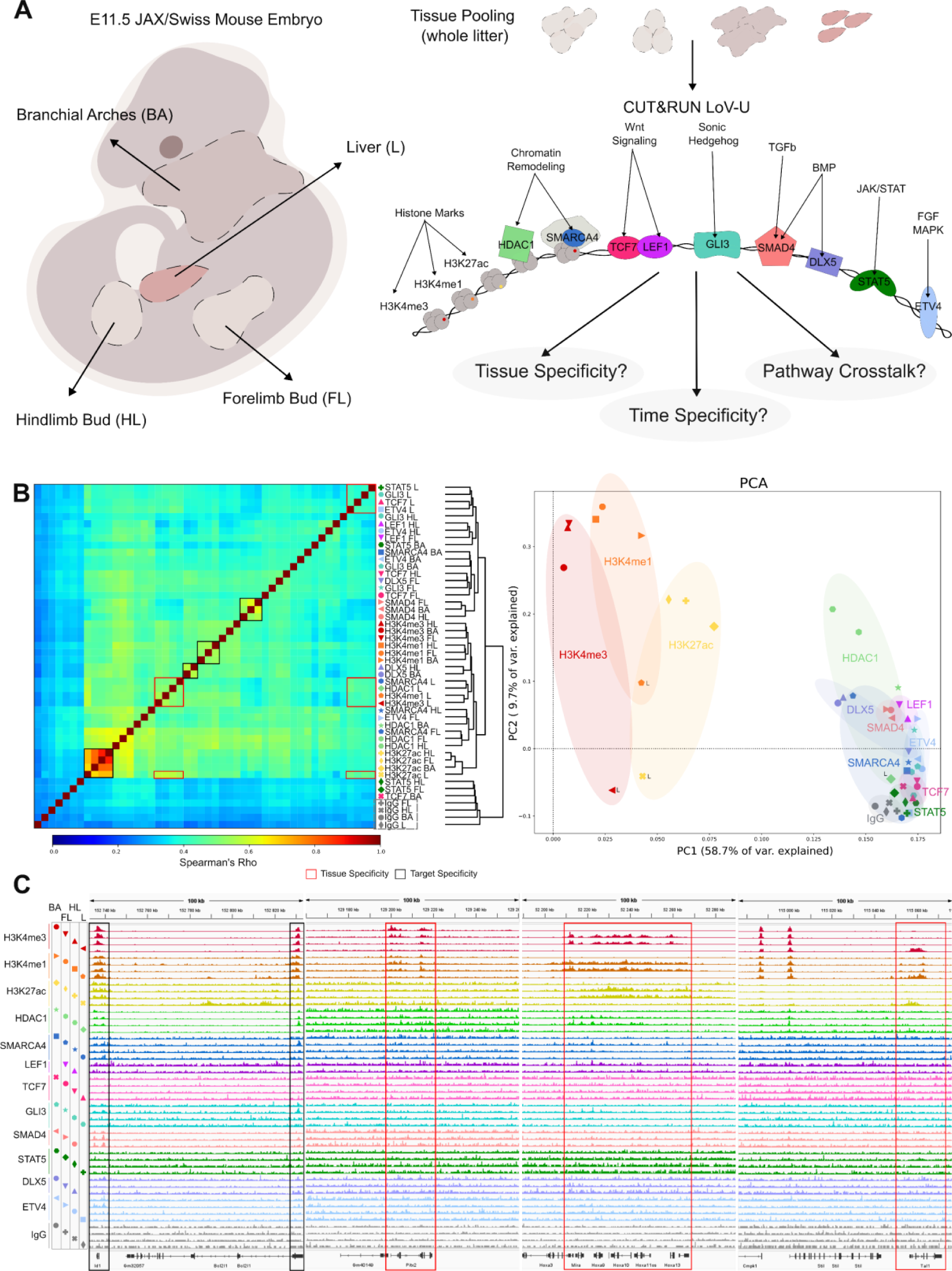
TFbomics of the E11.5 mouse embryo. **A.** Schematic overview of the experiment design, detailing the tissues dissected from E11.5 JAX/Swiss mice and the targets profiled in each tissue using C&R LoV-U. **B.** Left: Clustering of data according to Spearman’s correlation. Black boxes highlight higher correlation or clustering according to the profiled factor, whereas red boxes highlight this between factors within the same tissue. IgG negative controls are highlighted in a grey box. Right: Overall data patterns shown via PCA. PC1 (58.7% of variance) separated some samples by the target factor (ellipses shaded by factor color code), while PC2 (9.7% of variance) separated based on tissue (liver samples located towards negative values). **C.** Example tracks in loci showing little tissue specificity (far left), highest signal in BA (center left), highest signal in limb (center right) and highest signal in liver (far right). Tracks come from merged biological replicates normalized to reads per genome coverage and are group scaled by factor. Black boxes show factor specificity, red boxes tissue specificity. BA = branchial arches, FL = forelimb, HL = hindlimb, L = liver.

Crucial for achieving experimental success in C&R is the functionality of the antibody employed. We tested a collection of antibodies (often multiple per target) against >50 known developmental GRs. On average, 1 in 4 antibodies yielded reliable genome-wide signal, underlying the intrinsic difficulty of detecting lowly abundant nuclear proteins in their native conditions in C&R using commercially available antibodies. Here we present a curated set of genome-wide binding profiles (or tracks) of GRs that surpassed our experimental and computational quality controls across the four selected tissues (Figure 1B). The profiled targets include transcription factors and co-factors chosen to monitor the activity of key developmental pathways [ETV4 tracking Fgf signaling (Mao et al., 2009); STAT5A tracking jak/Stat signaling (Smith et al., 2023); DLX5 for Bmp signaling (Holleville et al., 2003); SMAD4 for Tgfb signaling (Guglielmi et al., 2021); GLI3 for Shh signaling (Büscher et al., 1997); TCF7 and LEF1 for Wnt signaling (Valenta et al., 2012)], chromatin regulators such as the histone deacetylase enzyme HDAC1 (Brunmeir et al., 2009) and the SWI/SNF component SMARCA4/BRG1 (Hota et al., 2022), and post-translational modifications on histones associated with active transcription such as H3K4me3, that signals active promoters (Schneider et al., 2004), H3K4me1 for active and poised enhancers (Bae and Lesch, 2020); and H3K27ac that marks promoters, enhancers, and super-enhancers (Calo and Wysocka, 2013). Producing over 100 individual C&R tracks allowed us to assess i) the tissue-specific genomic activity of developmental signaling pathways along with the resident chromatin dynamics, ii) the existence of convergent regulation or crosstalk of pathways in the same tissue, and iii) idiosyncrasies in the behavior of GRs (Figure 1A). The analyses to measure (i), (ii) and (iii) represent the bulk of this study.

Spearman’s correlation analyses followed by hierarchical clustering across all the datasets immediately revealed interesting patterns (Figure 1B with merged replicates, single replicates in Figure S1A). For example, we found groups of tracks that clustered based on one individual target, indicating instances where that target possessed a relatively similar binding activity across tissues, as for H3K27ac, H3K4me1 and – perhaps less expected – SMAD4 and DLX5 (black boxes in Figure 1B). In other cases, clustering was observed within tissues, where several factors appeared to co-regulate the same groups of genes in the liver (red boxes in Figure 1B). Notably, these groups were also marked by H3K27ac, likely reflecting the permissive chromatin state of the underlying loci (red flat boxes in Figure 1B). IgG negative controls clustered aside from all biological targets (grey box in Figure 1B), conferring overall credibility to the patterns of peaks identified. Consistently, Principal Component Analysis (PCA) separated the samples by factor along PC1, which explained ca. 58.7% of the variance (Figure 1B, right; ellipses shaded by factor color code, individual replicate PCA in Figure S1B), and by tissue along PC2, accounting for ca. 9.7% of the variance (Figure 1B, right; liver marked by L). Both patterns – overlap of GRs’ activity withing each tissue or in similarity of GRs’ binding profile across tissues – were visible upon close inspection of our tracks across exemplifying genomic loci: while several loci presented relatively homogenous signal across tissues (Figure 1C, black boxes on the left, tracks shown of merged biological replicates), many others displayed a high degree of tissue-specificity. These included binding sites within the locus of the neural crest regulator Pitx2 mostly and expectedly occurring in BA, the tissue containing cells from the neural crest (Kioussi et al., 2002) (Figure 1C, center left), the Hoxa gene cluster being prominently regulated in limbs (Nemec et al., 2017) (Figure 1C, center right), and the hematopoietic master gene Scl/Tal1 in L, the embryonic hematopoietic niche (Real et al., 2012) (Figure 1C, far right). Individual datasets at additional loci are shown in Figure S1C. The expected tissue-specific regulation of representative loci comforted us on the reliability of our experimental setup and consequent analytical output.

We compared our results with publicly available datasets. Of all the C&R peaks called across our analyses (58601 total unique regions), approximately half (44%; 25,820 total peaks) were called as open chromatin peaks in at least one tissue by the ENCODE ATAC-seq data which charted open chromatin regions in the limb, face and liver at E11.5 (Gorkin et al., 2020) (Figure S2A, left and center). Note that however, the other half of peaks (56%; 32,781 total peaks) presented higher-than-average ATAC-seq signal even if they were not called as open chromatin regions, and conversely many C&R datasets not called as peaks by us showed residual signal in ATAC-seq open chromatin regions (Figure S2A, signal intensity plots on the right). We also compared our C&R data with previously published ChIP-seq on H3K4me3, H3K4me1, and H3K27ac (Limb, facial prominence and liver E11.5; Gorkin et al., 2020) (Figure S2B-C), HDAC1 (Forelimb E11.5; Lex et al., 2020), SMARCA4 (Forelimb E11.5; Attanasio et al., 2014), GLI3 (C&R forelimb E9.5 and E10.5; Lex et al., 2020), SMAD4 (Forelimb E9.5 and E10.5; Gamart et al., 2021), DLX5 (Cerebellum; Holleville et al., 2003), STAT5 (Dendritic cells; Lee et al., 2022), ETV4 (EpRas cells; Arase et al., 2017), TCF7 and LEF1 (Nephron progenitor cells E16.5; Guo et al., 2021) (Figure S3) all conducted in different but comparable tissues and stages. Consistently with previous literature (Hu et al., 2023), C&R generally identified fewer peaks than the corresponding ChIP-seq data, but with a high degree of overlap (that is, most C&R peaks were called in the ChIP-seq). A lower concordance was observed for datasets which were poorly matched in terms of tissue or timepoint. Our matched datasets showed comparable signal profiles to ChIP-seq yet most often with a lower intensity, and generally contained signal in most regions called as peak in ChIP-seq (See Figure S2 for histone modifications and Figure S3 for other GRs). Different technologies and peak calling strategies might explain these differences, and typically ChIP-seq leads to more artifacts (Boyd et al., 2021; Hu et al., 2023). However, it is worthwhile to consider that thanks to using C&R we started each experiment from the equivalent material extracted from 3 limbs per sample, which is roughly 1/100th of the material required in a typical ChIP-seq (Zambanini et al., 2022; Zimmerli et al., 2020), granting our approach increased scalability, decreased material costs, and enormously improved ethical considerations.

### Tissue specific binding of gene regulators

GRs have pleiotropic activity in different tissues and their compound interactions are central to development (Andrey et al., 2017). We therefore decided to measure the differences and overlaps of the genome-wide binding across tissues for each targeted factor. We calculated how many peaks were identified in only 1, 2, 3 or across all 4 tissues, and plotted the resulting “decay” curves (Figure 2A). Some targets, such as H3K4me3 and H3K4me1, presented a relatively low slope with a great fraction of peaks still shared among 2, 3 and 4 tissues (Figure 2A, top left). This indicates a large overlap of active regulatory regions in the different developmental contexts. Most other targets had steeper slopes – the number of peaks was highest in a single tissue – hinting to higher tissue-specific actions (Figure 2A). For downstream analyses we focused on one example of a high-overlap/low decay (H3K4me3), one with intermediate slope (HDAC1) and one presenting a steep slope (GLI3) (Figure 2A).

**Figure 2.**
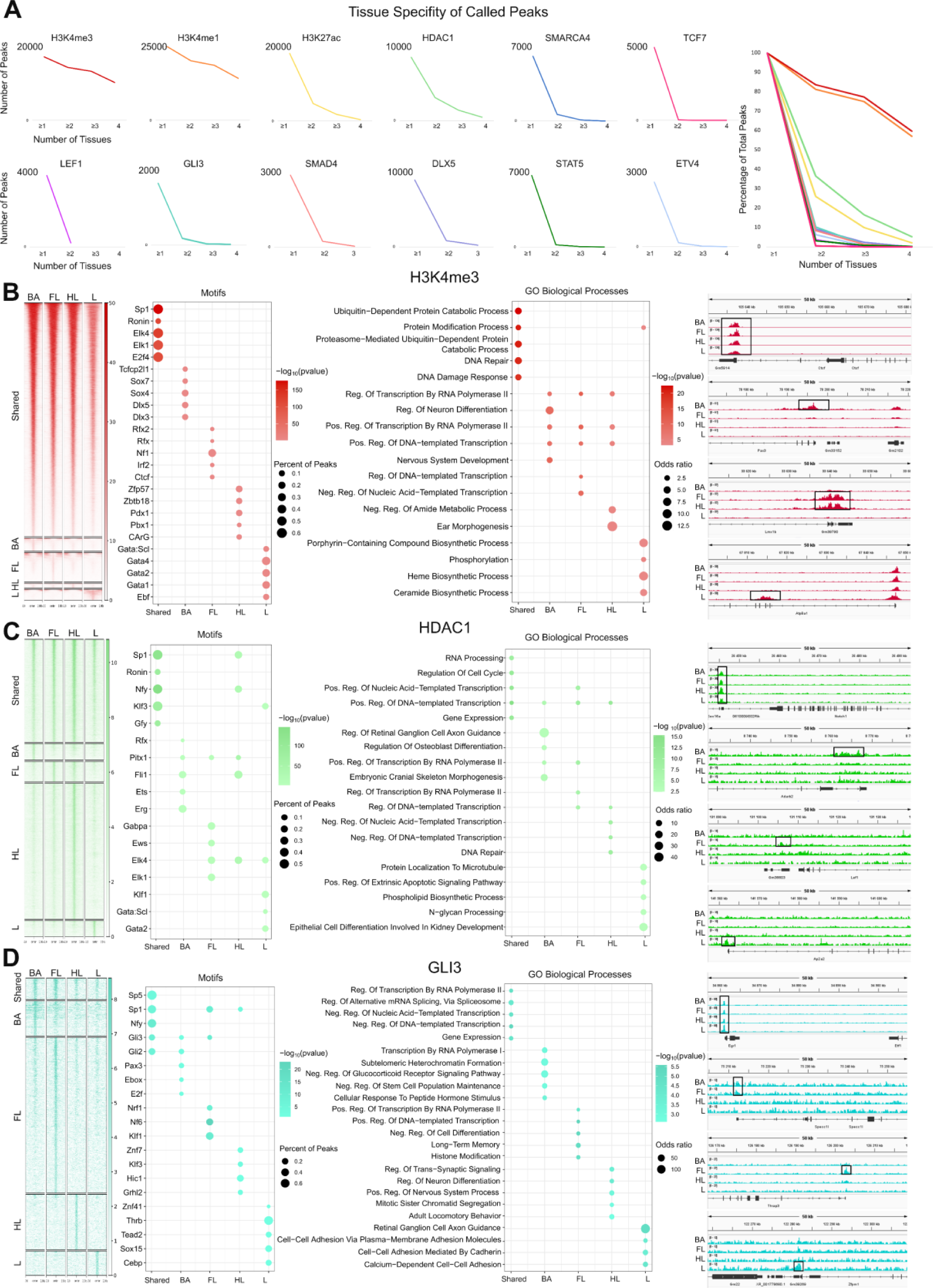
Tissue specificity of gene regulators. A. Decay plots showing number of peaks (y-axis) called per number of tissues (x-axis) for each factor. Right: Graph showing scaled decay curves for all factors in percentage of total peaks. B. Far left: signal profiles of H3K4me3 signal in shared, and tissue specific regions. Center left: enriched motifs in shared and tissue specific regions. Center right: enriched gene ontology biological processes of shared and tissue specific regions. Far right: example tracks showing shared and tissue specific regions. C. Signal profiles (far left), motifs (center left), gene ontology (center right) and example tracks (far right) for HDAC1 shared and tissue specific regions. D. Signal profiles (far left), motifs (center left), gene ontology (center right) and example tracks (far right) for GLI3 shared and tissue specific regions. Shared regions were defined as those called in 2 or more tissues. For motifs and gene ontology, color represents significance (p-value) and bubble size fraction of peaks (motifs) or odds ratio (gene ontology). The top 5 results (motifs and GO terms) for each peak set are shown. No q-value cutoff was applied. Tracks are a merge of two biological replicates, normalized with reads per genome coverage, and scaled by factor. BA = branchial arches, FL = forelimb, HL = hindlimb, L = liver.

The high overlap of H3K4me3-marked regions was perhaps surprising, as H3K4me3 decorates active and poised promoters (Schneider et al., 2004), a proxy of the transcribed genes underlying the differential gene expression programs, prerogative of organ formation. The large subset of shared regions might represent housekeeping genes, necessary for the function and survival of all cell types. We computationally separated the H3K4me3 peaks based on the signal being shared in at least two tissues or present only in one (Figure 2B). We found differential enrichment of TF binding motifs among these five groups (Figure 2B, left). The largest group of shared peaks presented motifs of SP and E2F, which are factors that are typically associated with general metabolic and cell-cycle related processes required by many cell types (Archer, 2011; Attwooll et al., 2004). The organ-specific H3K4me3 peaks, conversely, displayed motifs of known tissue-specific factors, such as for example SOX within BA (Schock and LaBonne, 2020) or GATA in the hematopoietic L (Jagannathan-Bogdan and Zon, 2013), among several others (Figure 2B, left). We annotated H3K4me3 peaks to genes and conducted gene ontology (GO) enrichment analysis for biological processes fostered by the shared and the non-shared H3K4me3-marked (active) genes. Shared genes belonged to ubiquitous processes such as protein processing and DNA repair, while the tissue-specific instances were enriched with other processes, including nervous system development in BA or heme biosynthesis in L (Figure 2B center, and example tracks on the right). Conceptual surprises, such as the presence of “ear morphogenesis” GO term in BA, implied ample room for discovery across our datasets, and likely represented a degree of pleiotropy determined by genes that can modulate more than one process.

The number of shared peaks in the HDAC1 dataset was proportionally smaller (Figure 2C, signal-intensity plot, left) and the examples of tissue-specific regulation were more numerous (Figure 2C left, and example tracks on the right). Similarly to H3K4me3, the sequences underlying unique HDAC1 peaks were also enriched in signatures of tissue-specific factors including again GATA and SCL in the L (Figure 2C). Interestingly, FL, HL and BA all presented motifs for PITX1 binding in non-overlapping peaks (Figure 2C, motifs on the left). PITX1 has been shown to be a HL-specific factor, acting on a developmental program shared between FL and HL (Nemec et al., 2017). Our data suggest that PITX1 is not limited to the core regulation of limb formation, but is active in other tissues, such as the BA, where it binds to different regulatory regions (again, the peaks presenting PITX1 motif are non-shared across tissues).

GLI3’s activity seemed to be predominantly tissue-specific with only few instances of peaks shared by at least two tissues (Figure 2D left, and example tracks on the right). As GLI3 is a DNA-binding protein we could confirm the presence of its cognate motif in the DNA sequence underlying the identified peaks (Figure 2D, motifs on the left). GLI3 motifs were highly enriched in BA and FL, emphasizing its function in these tissues (Akiyama et al., 2015; Elliott et al., 2020). Interestingly, while GLI3 drives digit number specification both in FL and HL (Lopez-Rios et al., 2012), our data suggest that it might do so by different mechanisms. The absence of GLI3 motif enrichment in HL and L points to its ability to cooperate with other DNA-binding transcription factors mainly as a cofactor (Figure 2D, motifs on the left), a bimodal modality of action similar to that which we previously observed for TBX3 (Amaia Jauregi-Miguel et al., 2023). These differential modes of action of GLI3 could provide a new rationale to mechanistically understand the multitude of human phenotypes caused by its mutations (Démurger et al., 2015).

### Binding of Gene Regulators Over Time

Gene regulatory networks are often dynamic and change along developmental trajectories (Tsankov et al., 2015; Zuniga and Zeller, 2020). We investigated the dynamics of GR binding by comparing two developmental time points in the liver - the site of embryonic hematopoiesis. We mapped the genome-wide binding profile of all the targets as above at the additional earlier time-point 10.5 dpc, to then compare it with the dataset obtained at 11.5 dpc (Figure 3A). Comparison between 10.5 and 11.5 dpc identified that TFs and CoFs tend to cluster within time-points (Figure 3B, left, red box), while chromatin marks are grouped depending on type of modification, regardless of time (Figure 3B, right, black boxes). We considered this interesting: the common localization of GRs downstream of diverse signaling pathways at a specific developmental stage might reflect their convergence onto broad genomic regulatory regions such as, as previously reported, super-enhancers (Hnisz et al., 2015). Our data indicate that these regulatory regions coregulated by GRs are different depending on the developmental time. Post-translational modifications of histones, on the other hand, seemed more stable over time, possibly indicating the consolidation of genomic identity that underlies the acquisition of a defined cell fate. Signal intensity profiles across all peaks supported this interpretation by showing how well intensity is conserved between 10.5 and 11.5 dpc in the case of H3K4me3 and H3K4me1, whilst it varies for other factors, in particular GLI3 and STAT5 (Figure 3C). Fluctuating signal over genomic regions, we reasoned, might reflect how dynamically these factors “jump” to new loci for the regulation of ontogenetically subsequent gene expression programs.

**Figure 3.**
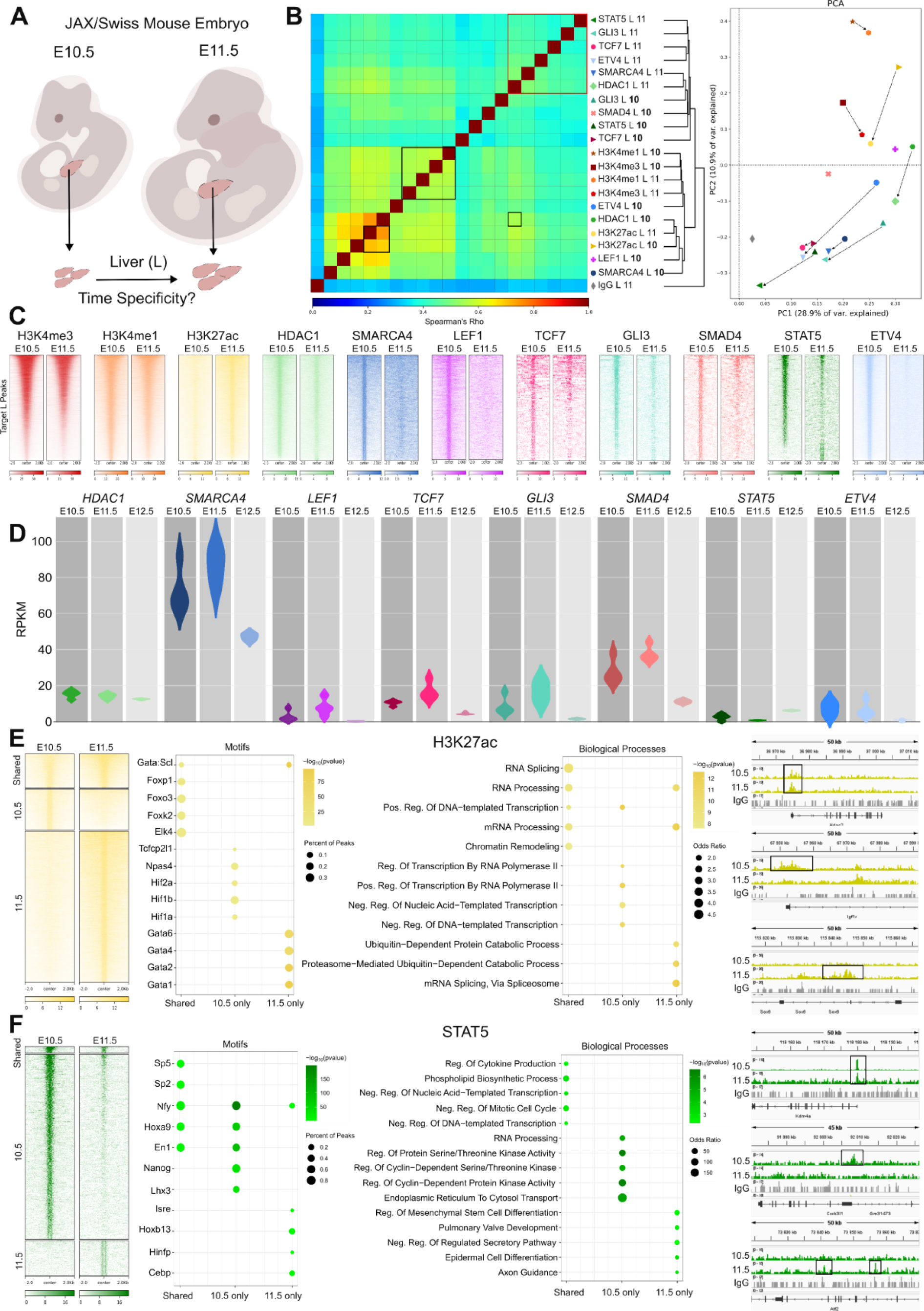
Time specificity of gene regulators. A. Schematic overview of the experiment design. B. Left: overall data patterns shown via Spearman’s correlation, showing correlation both by factor and time point. Right: Overall data patterns shown via PCA. Dashed arrows show connections between the same target over time. C. Heatmaps showing signal intensity for each target within all peak regions called at either time point for the specific target. LEF1 and SMAD4 11.5 datasets did not meet quality control standards and were thus excluded from earlier analyses, but are included here for signal comparison. D. Violin plots of RPKM values of RNA-seq data for genes coding for C&R target proteins at E10.5, E11.5 and E12.5 timepoints. N = 4, data from Cardoso-Moreira et al., 2019. E. Signal profiles (far left), motifs (center left), gene ontology (center right), and example tracks (far right) for H3K27ac shared and time specific regions. F. Signal profiles (far left), motifs (center left), gene ontology (center right) and example tracks (far right) for STAT5 shared and time specific regions. Shared regions were defined as those called in both time points. For motifs and gene ontology, color represents significance (p-value) and bubble size fraction of peaks (motifs) or odds ratio (gene ontology). Top 5 results for each peak set are shown. No q-value cutoff was applied. Tracks are a merge of two biological replicates, normalized with reads per genome coverage, and scaled by factor. L = liver.

The E11.5 datasets for LEF1 and SMAD4 were not included in the previous analysis as they failed our quality control metrics, and we never obtained successful DLX5 binding profiles in the liver at either timepoint. Multiple experimental attempts to obtain binding patterns of LEF1 and SMAD4 at E11.5 failed, despite that they were easily obtained at E10.5, and other targets worked well at 11.5 in parallel. These targets, in published RNA-seq (Cardoso-Moreira et al., 2019) exhibit an increase in expression at 10.5 followed by a drop in expression at 12.5 (Figure 3D). It is therefore plausible that these GRs are detectable only at an earlier time point.

We focused on H3K27ac (Figure 3E) and the seemingly dynamic regulator of embryonic hematopoiesis STAT5 (Schmerer et al., 2006) (Figure 3F) to provide an example of the type of analysis that our dataset allows. We divided peaks into three groups: i) shared between stages, ii) present in 10.5-only or ii) 11.5-only (Figure 3E, signal intensity plot on the left and example tracks on the right). In the case of H3K27ac-decorated regions, they displayed differential underlying transcription factors motifs: shared peaks were enriched in FOXO motifs, reflecting the role of FOXO factors in the establishment and maintenance of homeostatic stem cells (Menon and Ghaffari, 2018). 10.5-only peaks showed HIF1 consensus sequences, possibly reflecting the regulatory role of this TF in hematopoietic progenitor in hypoxic sites of the mouse embryo (Imanirad et al., 2014). Finally, 11.5-only peaks presented GATA motifs, most likely indicative of the prominent regulation of the fetal-to-adult switch of globin genes expression by GATA1, which starts in erythroblasts precisely from 11.5 dpc (Barbarani et al., 2021). We also ranked STAT5 peaks into three groups by the same logic (Figure 3F, signal intensity plot on the left and example tracks on the right). Interestingly, we found HOXA9 motifs in both the shared and 10.5-only STAT5 peaks. This is consistent with a role of STAT5 in early hematopoiesis (Schmerer et al., 2006) and with previous ChIP-Seq experiments conducted in HOXA9-transformed hematopoietic cells showing that STAT5 colocalizes with HOXA9 on regulatory regions (Ma et al., 2016). At 11.5-only we found HOXB13 instead of HOXA9 motifs despite – to the best of our knowledge – no function for HOXB13 in embryonic hematopoiesis has been described. HOX13 can regulate cell fate through association to super enhancers (Aspuria et al., 2020). It is plausible that STAT5 interplays with HOXB13 at later developmental stages or during adult hematopoiesis. Of note, at all stages, STAT5 seems to overlap with the regulator of hematopoietic proliferation and survival NF-Y (Bungartz et al., 2012). NF-Y was shown to regulate the expression of multiple *HOX* gene paralogs in hematopoietic stem cells (Zhu et al., 2005). Taken together, our data suggest the existence of a new gene regulatory network between STAT5, NF-Y and their regulation of – together with a possible physical interplay with – several HOX TFs.

### Genomic crosstalk between signaling pathways

Measuring how cell-communication signals are integrated on the genome is fundamental to understand cell differentiation during organ formation (Reményi et al., 2004). Our setup – that captures multiple transcriptional effectors of extracellular signals – allowed us to determine the interactions of these signaling pathways at the chromatin level. Here we focused on the FL, as a great wealth of data concerning the underlying gene regulatory networks exists for this tissue (Andrey et al., 2017; Zuniga and Zeller, 2020). Spearman’s correlation of all mapped factors at 11.5 dpc in FL unearthed several interesting observations. For example, we considered notable the proximity, within the clustering, between the chromatin remodelers SMARCA4, HDAC1 and H3K27ac-marked regions (Figure 4A). SMARCA4 (also known as BRG1), is part of the SWI/SNF complex that hydrolyzes ATP to remodel chromatin through nucleosome sliding and eviction, and it is typically associated with rendering the chromatin permissive for transcription (Centore et al., 2020). HDAC1 on the other hand, by removing acetyl groups deposited by CBP/p300 on H3K27 residues, represses transcription and prevents premature expression of developmental genes (Dsilva et al., 2023; Wang et al., 2022). Hence, the colocalization of HDAC1 and SMARCA4 on H3K27ac-positive regions was surprising. However, others have found that HDAC1 can be recruited within super-enhancers, that display H3K27ac, to activate transcription (Lai et al., 2023), or that the balance of p300 and HDAC1 activities controls the nucleosome eviction by the BRG1/SMARCA4-containing SWI/SNF complex in macrophages (Pietrzak et al., 2019). Moreover, our recent work in the context of Wnt signaling implied HDAC inhibition in the inability to open the chromatin for transcription (Pagella et al., 2023) which could be interpreted as a function of HDAC enzymes in gene activation rather than repression. The tight association between these factors in FL cells might indicate that, in this context, SMARCA4 and HDAC1 interplay on maintaining the chromatin open. Accordingly, most of the overlapping peaks between SMARCA4 and HDAC1 are either on H3K27ac or H3K4me1/me3 positive regions, which are both associated with active and transcription-permissive chromatin (Figure 4B). Concerning HDAC1, while its role in limb organogenesis has not been addressed, our data indicate a requirement of this factor for correct FL morphogenesis on a molecular level, complementing other phenotypic reports (Paradis and Hales, 2015).

**Figure 4.**
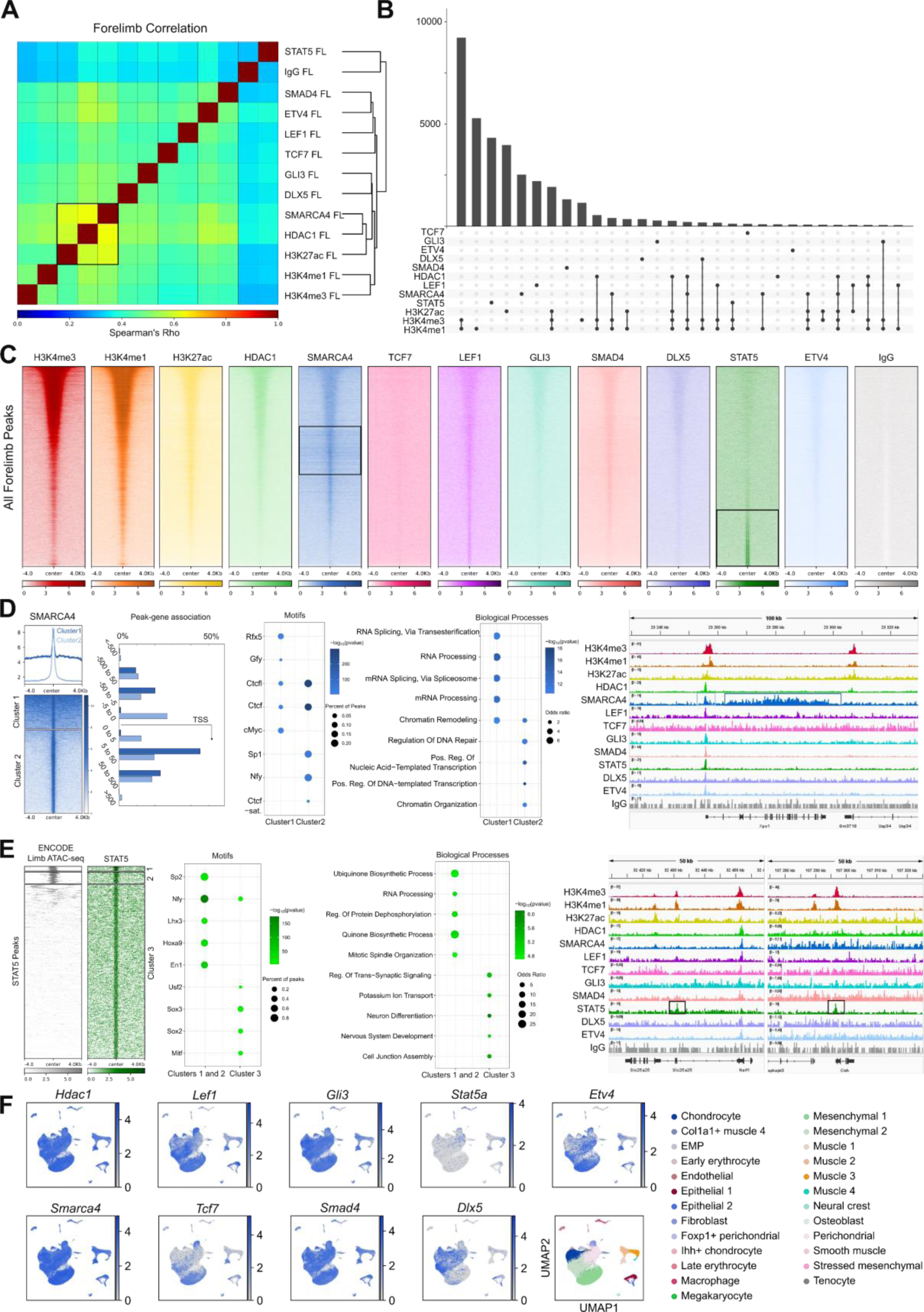
Crosstalk between signaling pathways. A. Overall data patterns shown by Spearman’s correlation. H3K27ac correlated with the chromatin modulators Smarca4 and Hdac1. Stat5 was separated from the others. B. Upset plot showing peak overlaps. The highest overlaps were between histone modification datasets. C. Heatmaps showing signal intensity for all targets across all the peaks called in the forelimb. Smarca4 and Stat5 emerged as having unique signal profiles (black boxes). D. Far left: clustering of Smarca4 revealed two modes of enrichment. They differed in their proximity to genes and in the average expression of associated genes. Motifs (center left), gene ontology (center right) and example tracks (far right) exemplify the differences between the two Smarca4 peak clusters. E. Far left: ENCODE ATAC-seq signal in the limb at E11.5 plotted at Stat5 peaks revealed that most Stat5 peaks were defined by closed chromatin. Motifs (center left), gene ontology (center right) and example tracks (far right) show differences in the Stat5 peaks within open (clusters 1 and 2) and closed (cluster 3) chromatin. Tracks on the right show two instances of cluster 3 peaks, where Stat5 is enriched in regions that lack signal from the other profiled transcription factors. F. scRNA-seq data from He et al., 2020 showing cell-type specific expression patterns of genes encoding for C&R targets. FL = forelimb. For motifs and gene ontology, color represents significance (p-value) and bubble size fraction of peaks (motifs) or odds ratio (gene ontology). Top 5 results for each peak set are shown. No q-value cutoff was applied. Tracks are a merge of two biological replicates, normalized with reads per genome coverage, and scaled by factor.

We noticed another peculiarity when visually inspecting the binding pattern of SMARCA4. When plotting signal intensity profiles of all genomic loci in which we found a peak for at least one factor, we noticed that a fraction of SMARCA4 peaks displays broad (instead of sharp) signal (squared section in the SMARCA4 plot, Figure 4C). SMARCA4 binding can indeed be separated into two distinctive behaviors, with peaks surrounded by high signal or broad domains (cluster 1 in Figure 4D) and sharp peaks (cluster 2 in Figure 4D). Both clusters are represented by numerous instances (that is, neither of these two is a rare behavior; Figure 4D, bottom). Notably, cluster2/broad domains marked regions upstream and downstream of the transcriptional start site (TSS), often decorating the entirety of a gene body, while cluster1/sharp peaks are most often found downstream of the TSS (Figure 2D, peak-gene association, and example locus on the right). This pattern of SMARCA4 binding, though absent in the published ChIP-seq that we used as comparator, was confirmed via C&R in HEK293T cells (Figure S3F and S3G). Cluster 1 and 2 are characterized by the presence of different transcription factors motifs, and the annotated genes belong to different GO categories. This suggests that differential cooperativity between SMARCA4 and various transcriptional complexes is required to execute separate functions, such as chromatin remodeling (cluster 1) or DNA repair (cluster 2) (Figure 4D, see Motifs and GO Biological Processes).

Finally, we noticed that STAT5 clusters separately from all other datasets, similarly to the IgG control (Figure 4A). This could not be explained by poor signal: on the contrary, STAT5 dataset was characterized by high signal-to-noise ratio (Figure 4E, note the green signal intensity plot). Most of STAT5 peaks did not overlap with any other dataset, except for H3K4me1/3-positive regions (Figure 4B). This behavior of STAT5 as a ‘lone wolf’ was also apparent when signal intensities across all genomic loci through which we identified binding for at least one factor were compared. Here, STAT5 clustered on the bottom in regions where the average signal of all other factors, was low (Figure 4E, boxed loci in the STAT5 chart; also see example track on the right where STAT5 binds on positions lacking signal from the other factors). This surprised us, as STAT5 has been found to mediate chromatin interactions in super-enhancers of immune cells (Hennighausen and Lee, 2020; Li et al., 2017). Studies of STAT5 in embryological contexts such as limb formation are lacking, but our dataset supports that its function might be mechanistically different from that in the adult immune system – an observation that warrants more work on the developmental roles of STAT5. Notably, comparison of STAT5 peaks to open chromatin measured via ATAC-seq in FL at E11.5 dpc revealed that most STAT5 peaks were found in closed chromatin. STAT5 peaks in closed chromatin (cluster 3 in Figure 4E) presented motifs of SOX factors (center left), suggesting the tantalizing possibility that STAT5 plays a role in the SOX-mediated chromatin pioneering (Julian et al., 2017). This confirms the recent identification that STAT5 is the earliest factor binding and remodeling the *Il9* locus in human T-helper cells (Fu et al., 2020), and underlies a novel mechanism of the reprogramming of cells within the limb bud that will acquire novel differentiation fates.

One limitation of our bulk approach is that it does not allow to discern instances of locus-coregulation by GRs in the same cell, from different factors being active on the same regulatory regions but in different cells. To disentangle this, we used existing single-cell RNA sequencing data (He et al., 2020) and plotted the cell-type specific expression of each of our C&R targets (Figure 4F). While some genes, such as *Hdac1* and *Smarca4*, had a broad expression pattern encompassing many cell types, certain targets showed more of a tissue specific distribution. For example, *Tcf7* and *Lef1* expression was highest in the different mesenchyme and fibroblast clusters, respectively, indicating that our TCF7/LEF1-specific C&R reads should be traced to these cell types (Figure 4F). *Stat5a* was lowly expressed overall but had detectable expression in macrophages, consistent with its known role in the immune system, as well as in chondrocytes and specific types of muscle cells (Figure 4F). Therefore, we conclude that STAT5A regulates the identified targets in this cell type. This simple data integration shows that single-cell RNA-seq datasets – and by extension also spatial transcriptomics results – can be proficiently used to extrapolate from which cell population the C&R signal derives.

### Convergence of genomic regulators

Our FL tracks combined identified 36717 peaks called in at least one dataset – FL-specific regulatory regions – many of which (ca. 18000) are bound by only one of the factors that we tested (Figure 5A). We generated lists of peaks that were bound by more than one with at least 2, 3, 4 – and so on – factors and noticed that the number of regulatory regions contacted by an increasing number of GRs drops rapidly (Figure 5A). This possibly underscores that combinations of GRs and DNA regulatory elements are the most common means to drive the expression of differential gene expression programs (as opposed, for example, to having different GRs binding on the same regulatory element or the same GR binding on different regulatory elements). We were surprised, however, in noticing that the groups of peaks presenting a higher number of bound GRs – to which we colloquially refer to as “popular” regulatory regions – were characterized by increased evolutionary conservation as assessed with phastCons, which measures the degree of negative selection of a region on a scale of 0 (no conservation) to 1 (full conservation) (Figure 5B). The trend became statistically significant when we divided all regulatory regions in 4 groups: called in 1-2 datasets, 3-5, 6-8, and 9-12, indicating that the regions called as peaks in a greater number of datasets are evolutionarily conserved to a higher degree and thus likely essential for gene regulation (Figure 5C). We focused on the group with the highest statistical score, that included 99 peaks presenting the concurrent presence of at least 75% (9 out of 12) of our targets (H3K4me1; H3K4me3; H3K27ac; HDAC1; SMARCA4; LEF1; TCF7; GLI3; SMAD4; DLX5; STAT5; ETV4). All these 99 genomic regions presented high signal/noise for all factors, while signal was absent in the IgG control samples (Figure S4A). This group of genomic elements, we hypothesized, might represent an ensemble of important hubs where several pathways converge to regulate the expression of essential genes for limb patterning. Consistently with this, motif analysis revealed enrichment of known determinants of limb morphogenesis (Figure 5D), such as the LIM homeobox factor LHX9, which was shown to integrate signaling pathways to control the three-dimensional patterning of growing vertebrate limbs (Tzchori et al., 2009), and Engrailed-1 (EN1), one of the initial inducers of the apical ectodermal ridge (Loomis et al., 1996). The relevance of these regulatory regions seemed to go beyond limb patterning: STRING showed a high interconnection between the 99 hub regions and the targets themselves (Figure 5E), and GREAT annotated these 99 regulatory regions to genes that, when mutated, cause early embryonic lethality (Figure 5E, grey bubbles, obs/exp 3.24, FDR 0.028). Among these are the ubiquitously functional DNA repair protein BRCA1 (Hakem et al., 1998) and the histone methyltransferase PRMT1, whose absence leads to stalled development in vertebrate model systems (Shibata et al., 2020) (Figure 5F, example tracks). We looked to see how highly bound these regions were across tissues, and found that also in the HL, BA and L, these regions showed high signal in many targets (Figure S4B). Interestingly, when we plotted Spearman’s correlation of all the datasets within the hub regions, we found that target specific idiosyncrasies allowed for clustering that was based highly on the target protein (Figure S4B). To explore the possible connotations and functions of these regions within the human genome, we converted the coordinates from mm10 to hg38 (minimum remap ratio 0.5). Of the 99 regions, 88 successfully mapped onto hg38 – in comparison, of 99 regions randomly selected from all of the FL peaks, only 66 successfully converted. In humans, these regions generally belong to CpG islands and coincide with ENCODE candidate cis-regulatory elements, both enhancers (yellow) and promoters (red) (Figure 5G). A high level of ChIP-seq reMap signal in most regions indicated that, like in our mouse datasets, these are “popular” regions bound by a multitude of GRs (Figure 5G). JARVIS scores of negative selection in humans were also high for these regions, suggesting that they retain their functional relevance for human development.

**Figure 5.**
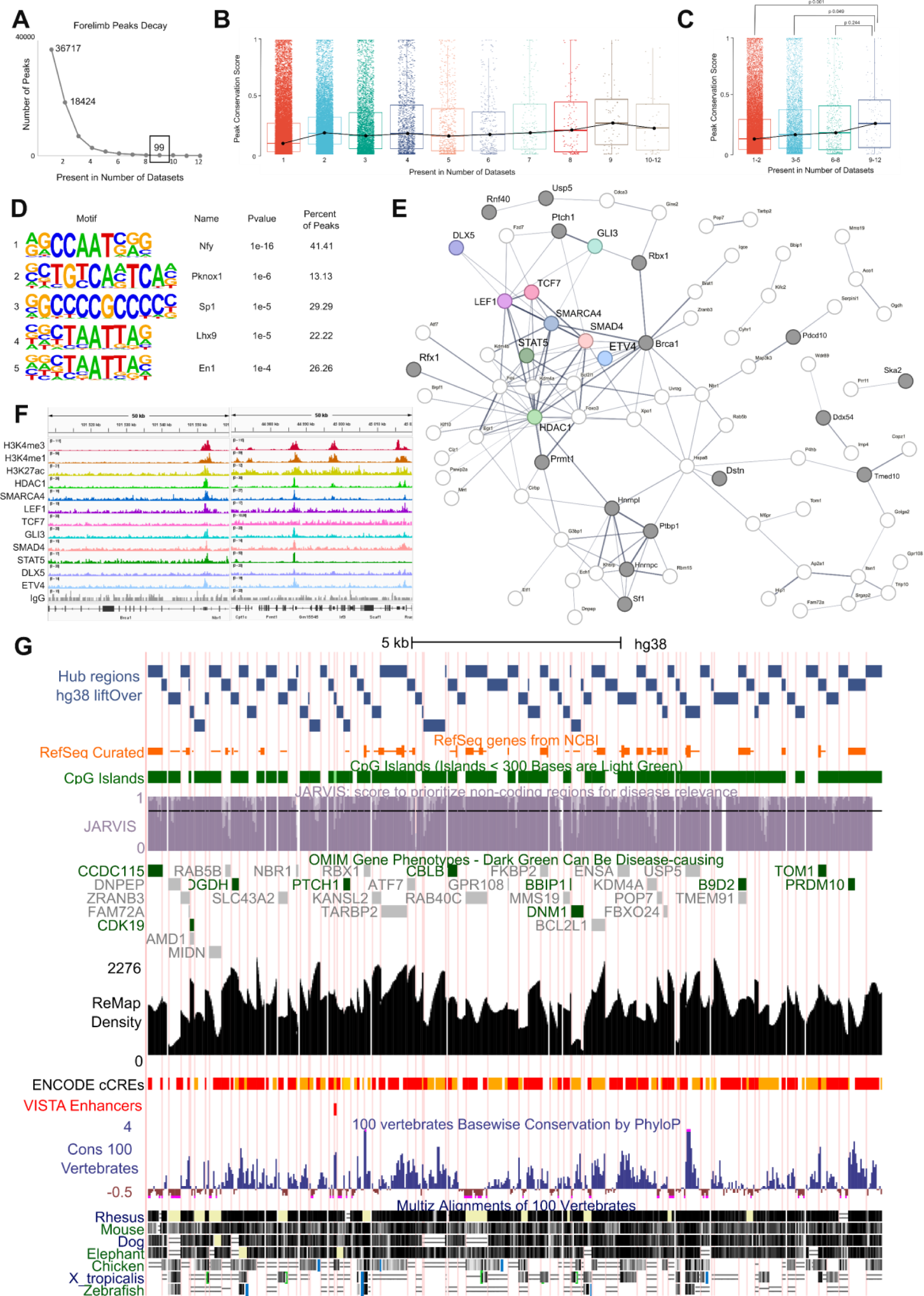
Identification of highly bound regulatory hubs. A. Graph showing the peak decay within the forelimb, on the y-axis is the number of peaks and on the x-axis the number of datasets in which they were identified. 99 regions were called in at least 9 of 12 datasets, these were defined as potential regulatory hubs. B. phastCons-based scoring of the 12 groups of peaks identified in A. Negative selection is measured on a scale from 0 (no conservation) to 1 (full conservation). C. The trend shown in B becomes statistically significant when the 12 groups are merged in 4: peaks called in 1-2 datasets, 3-5, 6-8, and 9-12. D. Top enriched motifs within the hub regions include promoter and high GC motifs, as well as developmental transcription factors. E. STRING map showing the 99 hub regions plus the targets themselves. The protein networks were highly interconnected, with enrichment for early embryonic lethality prior to organogenesis (grey bubbles; obs/exp 3.24, FDR 0.028). Line thickness indicates interaction confidence. Disconnected nodes were removed. F. Example tracks showing hub region loci. Tracks are a merge of two biological replicates, normalized with reads per genome coverage and scaled individually. G. UCSC genome browser view of the mm10 hub regions lifted over to the hg38 genome, showing RefSeq genes, CpG islands, JARVIS score of negative selection in human populations, OMIM gene phenotypes, ReMap ChIP-seq peak density, ENCODE candidate cis-regulatory elements, VISTA enhancers, and conservation values.

## Discussion

The central question of developmental biology is how the cells of an organism, despite having inherited the same genome, can build an embryo by activating the right genes at the right time to ensure the desired differentiation programs. Decades of research in molecular biology have clarified that this problem could be solved by determining the activity of transcription factors, cofactors, and other GRs, which bind the DNA at gene regulatory regions and contribute to gene activation or repression (Dickel et al., 2013; Dunham et al., 2012). Several technologies, including ChIP-seq and now the more proficient C&R, allow measuring where a given GR binds across the genome. However, ChIP-seq, C&R, and all the related technologies are limited to detecting one target at the time, as they rely on the use of protein-specific antibodies. This limitation has caused researchers to focus on one, or on a limited numbers of, favorite GR candidates. However, GRs do not act alone. Rather, they are needed in a combinatorial fashion to functionally engage regulatory regions and modulate gene transcription (Reményi et al., 2004). Hence, the study of gene expression will greatly benefit from the parallel measurement of all GRs acting in the context of interest.

Here, we initiated the assessment of the genome-wide binding profiles of multiple GRs in selected developmental contexts as an attempt to show the feasibility of a larger-scale venture. Certainly, the sheer number of GRs encoded in the vertebrate genome – ca. 1600 transcription factors and likely a comparable number of co-factors and chromatin remodelers – might be discouraging and render this endeavor unattainable. Big consortia, such as ENCODE, successfully approached this enormous task by mapping the genome-wide binding site of many GRs using ChIP-seq (Landt et al., 2012; Luo et al., 2020). However, most of this work is conducted in human cell lines (Partridge et al., 2020), for which it is not always clear to what extent they are representative of the tissue or disease of origin (Ben-David et al., 2018; Lopes-Ramos et al., 2017). This is particularly relevant when desiring to study physiological mechanisms occurring during normal embryonic development. With the current study, we started and support the next step: the monitoring of all GRs in relevant *in vivo* model of developmental processes: the resulting information concerning the combinatorial action of GRs could be used to reveal how the genome is deployed in different organs during development to generate cell-type specific gene expression programs. We advocate for the gradual construction of an Atlas describing the action of virtually every GR acting during different moments of organogenesis. This initiative, to which we refer as Transcription Factor Binding-Omics (TFBomics) will constitute a colossal repository whereby researchers interested in gene regulation during development, can access, download, or simply consult the data instead of conducting sophisticated experiments such as ChIP-seq and C&R.

Despite its limiting size, the current study is already enough to allow analytical possibilities that permit new discoveries, among the many that this atlas could offer. Here we will briefly mention three findings that we consider being the most relevant. Our first discovery derives from the exploration of how the activity of select GRs varies across tissues at the same developmental time (Figure 2). Several GRs possess ample tissue-specific activity – that is, peaks identified in one tissue and not in the others – but also common actions, whereby they appear to regulate the same DNA elements in different tissues. This provides a clear-cut molecular rationale for gene pleiotropy. It is interesting to note that, when the binding signal of a given GR is compared between tissues, we observed instances of variability in its intensity (see for example HDAC1 in Figure 2C) indicating that GRs are present but bind with lower affinity or in varying fractions of the cell population depending on the tissue, as well as more black-and-white cases (see for example the HL-only peaks of GLI3 in Figure 2D) where sets of tissue-specific targets seem to be exclusive of a particular developmental trajectory. Our second discovery is the investigation of how GRs act over time. Domcke and Shendure recently advocated for the construction of cell trees rather than atlases, as these will better explain functional gene expression during ontogeny (Domcke and Shendure, 2023). We profoundly agree, and have the ambition to make the TFBomics dataset grow and include as many organ-specific differentiation time-points as possible. Two stages of liver development, for example, were enough to discover that the two time-points (10.5 and 11.5 dpc), despite being relatively close, present a conspicuous rewiring of the chromatin landscape (see H3K27ac in Figure 3E). This constitutes rationale for raising the awareness that generalizing the binding pattern of a GRs to other models, tissues, and even time-points, when only one context has been assessed, might be incorrect, and provides the impetus to perform large scale profiling of GRs while considering time-course trajectories. The third novel observation, and perhaps the most thought-provoking one, concerns that identification of “popular” regulatory regions, frequented by a conspicuous number of GRs. These are lower in number than other regulatory regions bound by one or only a few GRs (Figure 5A). However, they present a statistically higher evolutionary conservation. Interestingly, the popular regulatory regions we identified were not necessarily super-enhancers (Figure 5), which were previously suggested being regulatory hubs where signaling cascade converge (Hnisz et al., 2015). We propose that different signaling cascade evolved to regulate distinct groups of genes depending on positional information whilst all cooperating to maintain the expression of cell-essential genes by acting on conserved genomic elements that can simultaneously accommodate a multitude of factors. One should keep in mind that a deep form of activity-conservation that goes beyond sequence conservation, but concerns a regulatory syntax where enhancers can be interpreted by the available GRs, has been shown to exist (Wong et al., 2020). Finally, whether these regions are crowded – that is, GRs bind simultaneously – of bound in different cell types or alternatively along shorter timescales, remain to be determined.

One main limitation currently exists in the construction of the TFBomics. C&R-LoV-U is a bulk-cell population approach. While single cell C&T exists (Bartosovic et al., 2021) and few reports of success are available, from extensive contacts with the community and experience in our laboratory we understand that most laboratories still recur to ChIP-seq for mapping dynamic transcription factors or non-DNA-binding co-factors, where both C&R and C&T are often unsuccessful. While ChIP-seq might still present advantages and result in a higher number of binding events given by the large number of cells used and the application of chemical cross-linking, we favor C&R and C&T as they allow the reduction of material needed of one or two orders of magnitude, and do not require cross-linking, thereby eliminating the artifacts typically introduced by this procedure. Moreover, our recent C&R-LoV-U protocol is successful with almost all GRs that we have tested so far, even when targeting dynamic, non-DNA binding proteins (Zambanini et al., 2022). Using C&R-LoV-U is, in our opinion, the current best choice for the large-scale parallel profiling of GRs. scRNAseq and spatial transcriptomics technologies can be synergistically combined with C&R to determine the precise expression of each GRs, thereby increasing the spatial resolution of the C&R findings.

## Acknowledgments

This work was supported by funding to C.C. from the Swedish Research Council, Vetenskapsrådet [2021-03075 and 2023-01898], Linköping University and LiU/RÖ-Cancer, Cancerfonden [CAN 2018/542 and 21 1572 Pj], and Additional Ventures (USA) [SVRF2021-1048003]. C.C. is a Wallenberg Molecular Medicine (WCMM) fellow and receives generous financial support from the Knut and Alice Wallenberg Foundation. The computations and data handling were enabled by resources provided by the National Supercomputer Centre (NSC), funded by Linköping University; Peter Münger at the National Supercomputer Centre is acknowledged for assistance concerning technical and implementational aspects in making the codes run on the Sigma resource. Funding for open access charge: Library of the Linköping University (Bibsam affiliated).

## Author contributions

A.N. and G.Z. performed CUT&RUN experiments and analyzed the data. A.N. integrated all the analyses and designed the figures. M.J. assisted with the initial experimental setup. Y.vdG. and T.W. integrated the scRNA-seq data. P.P. assisted with mouse tissue isolation and project design. C.C. conceived the project, provided the funding, supervised the research team, and wrote the manuscript. All the authors revised the manuscript and the figures.

## Methods

### Cell Culture

Human embryonic kidney 293T cells (HEK293T) were cultured at 37°C in a humidified incubator with 5% CO2 in high glucose Dulbecco’s Modified Eagle Medium (41965039, Gibco) supplemented with 10% bovine calf serum (1233C, Sigma-Aldrich) and 1× penicillin-streptomycin (15140148, Gibco).

### Animal experimentation

Animal housing and experimentations were performed under the ethical animal work license obtained by C.C. at Jordbruksverket (Dnr 2456-2019 and 03741-2024), and all procedures and housing were according to the Swedish laws and guidelines. JAX Swiss Outbred mice (strain 034608) were used. Housing was in Allentown NexGen IVCs, floor area 500 cm2 with a maximum of 4 mice/cage. Cages had aspen wood shavings and two types of shredded paper for nesting, and paper tubes were also provided. Temperature was 21±2°C, humidity 45-65% and light cycle 12 h/12 h (7.00 am/7.00 pm). Mice had unrestricted access to sterilized drinking water, and ad libitum access to a pelleted and extruded mouse diet. Mice were housed in a barrier-protected specific pathogen-free unit, and this status was monitored and confirmed according to FELASA guidelines by a sentinel program. The mice were free of all viral, bacterial and parasitic pathogens listed in FELASA recommendations (Mähler et al., 2014). Embryo age was determined based on timed mating and vaginal plug observation (0.5 dpc) and confirmed by morphological criteria. Experiments were performed in the afternoons (female sacrifice at approximately 13:00). Pregnant females were sacrificed by cervical dislocation and E10.5 or E11.5 embryos were surgically removed. Tissues were dissected under a dissection stereomicroscope (SZ61, Olympus) and pooled from each litter.

### CUT&RUN-LoV-U

Tissue pools were dissociated to single cells via incubation in TrypLE Express Enzyme (12-604-013, Thermo Fisher Scientific) for 15 min at 37°C on a shaker. For liver tissues, dissociation was done with manual pipetting. The cell suspension was resuspended in ice-cold PBS, filtered through a 40 µm cell strainer (KKE3.1, Carl Roth) and further processed for C&R. 3 limbs were used per sample, while the equivalent of 1 liver or 1 set of branchial arches were used per sample. For HEK293T cells, 500,000 cells were used per sample.

C&R LoV-U was performed as described in Zambanini et al., 2022. Filtered cells were washed three times in nuclear extraction (NE) buffer [20 mM HEPES-KOH (pH 8.2), 10 mM KCl, 0.5 mM spermidine, 0.05% IGEPAL, 20% glycerol, Roche Complete Protease Inhibitor EDTA-Free], the resuspended in 40 µl NE per sample and bound to 20 µl magnetic ConA agarose beads (ABIN6952467, antibodies-online) previously equilibrated in 44 ul binding buffer per sample. After incubation, nuclei and beads were resuspended for 5 min in EDTA wash buffer (wash buffer [20 mM HEPES (pH 7.5), 150 mM NaCl, 0.5 mM spermidine in Roche Complete Protease Inhibitor EDTA-Free (COEDTAFRO)] with 2 mM EDTA) and split into PCR tubes. Antibody incubation was performed in 200 µl wash buffer with antibody dilute 1:100 (for antibody and batch information, see Table S1) overnight at 4°C on a rotator. After overnight incubation, samples were washed five times in wash buffer and resuspended in 200 µl of pAG-MN buffer (wash buffer with pAG-MN 120 ng/sample) and incubated for 30 min at 4°C on a rotator. After 5 washes, digestion was performed for 30 min in ice: started by resuspending the samples in 50 µl of wash buffer with 2 mM CaCl2. After 30 min, the digestion buffer was removed and moved to tubes containing 3 µl EDTA/EGTA 250 mM. The digestion reaction on the beads was stopped with 50 µl of 1× Urea STOP buffer (100 mM NaCl, 2 mM EDTA, 2 mM EGTA, 0.5% IGEPAL, 8.5 M urea) and the samples were incubated 1 h at 4°C. Beads were collected on the magnet and liquid transferred to the PCR tube containing the digestion buffer. DNA was purified by two subsequent rounds of bead-based cleanup using Mag-Bind TotalPure NGS beads (M1327, Omega Bio-Tek) at 2×, and then resuspended in 20 µl Tris-HCl (pH 7.5).

### Library preparation and sequencing

Library preparation was performed with KAPA Hyper Prep Kit for Illumina platforms (KK8504, KAPA Biosystems) according to the manufacturer’s guidelines with modifications: End repair and A-tailing was performed in 0.4× volume reactions with 20 µl of purified DNA. The thermocycler conditions were set to 12°C for 15 min, 37°C for 15 min and 58°C for 25 min to prevent thermal degradation of the shortest fragments. Adapter ligation was performed in 0.4× volume reactions. KAPA Dual Indexed adapters were used at 0.15 µM. A post-ligation clean-up was performed with Mag-Bind TotalPure NGS beads at 1.2× the sample volume. Resuspension was carried out in 10 mM Tris-HCl (pH 8.0). Library amplification was performed in 0.5× volume reactions. The cycling conditions were set as follows: 45 s initial denaturation at 98°C, 15 s denaturation at 98°C, 10 s annealing/elongation at 60°C, 1 min final extension at 72°C, hold at 4°C, with 13 cycles. After amplification, a post-amplification clean-up was performed using NGS beads at 1.2× sample volume. Libraries were then run on an E-Gel EX 2% agarose gel (G402022, Invitrogen) for 10 min using the E-Gel Power Snap Electrophoresis System (Invitrogen). Bands of interest between 150 and 500 bp were excised and purified using the GeneJET Gel Extraction Kit (K0691, Thermo Fisher Scientific) according to manufacturer’s instructions. Libraries were quantified with the Qubit (Thermo Scientific) using the high sensitivity DNA kit (Q32854, Thermo Fisher Scientific), pooled and sequenced 36 bp pair-end on the NextSeq 550 (Illumina) using the Illumina NextSeq 500/550 High Output Kit v2.5 (75 cycles) (20024906, Illumina) to an approximate depth of 5-10 million reads per sample.

### Read trimming and mapping

Quality of reads was assessed using fastqc (Brandine and Smith, 2022, version 0.11.9). Raw reads were trimmed with bbmap bbduk (version 38.18, (Bushnell et al., 2017) to remove adapters, artifacts, [AT]18, [TA]18, and poly G or C repeats. Reads were aligned to the mm10 or hg38 genome with bowtie2 (version 2.4.5, (Langmead and Salzberg, 2012)) with the options –local –very-sensitive-local –no-unal –no-mixed -no-discordant –phred33 –dovetail -I 0 -X 400. SAMtools (version 1.11, (Li et al., 2009)) was used to create bam files, fix mate pairing, and for deduplication. Bam files were then filtered to remove reads falling within C&R suspect list regions (Nordin et al., 2023).

### Quality control

Individual replicates were assessed based on duplication rate after trimming (<10%), sequencing depth of mapped, deduplicated, and suspect list filtered bam files (typically 4 – 10 million reads), fragment profiles (TFs show mostly sub- and mono-nucleosome fragments, histone marks typically show mono- and di-nucleosomes), signal at positive control loci (known binding sites for the target), and concordance between biological replicates with genome wide binning, and/or peak calling.

### Peak calling and visualization

For peak calling, visualization and signal graphs, replicate bam files were merged with samtools into a single file. Bedgraphs were created with BEDtools (version 2.23.0, (Quinlan and Hall, 2010)) genomecov on pair-end mode and default settings. Peaks were called using MACS2 (Zhang et al., 2008, version 2.2.6) with the options -f BAMPE and -p 1e-3. deepTools (version 3.5.1-0, Ramírez et al., 2014) was used to convert bam files to normalized bigwig files (bamCoverage using -RPGC option and -e to extend reads to fragment length). Bigwigs were visualized in IGV, tracks shown in the figures are a merge of two biological replicates, normalized with reads per genome coverage, and scaled by factor.

### Downstream Analyses

Deeptools was used to make signal intensity heatmaps, correlation plots and PCAs. Peak overlaps were performed using BEDtools. Venn diagrams and Upset plots were created using Intervene (Khan and Mathelier, 2017, version 0.6.4). Peak-gene annotation was performed using GREAT (version 4.0.4, McLean et al., 2010) on default settings. Motif analysis was done with HOMER (version 4.11, Heinz et al., 2010) with the option -size given. GO analysis was done using Enrichr (Kuleshov et al., 2016). For motifs and gene ontology visualization, color represents significance (p-value) and bubble size fraction of peaks (motifs) or odds ratio (gene ontology). Top 5 results for each peak set are shown regardless of significance: no q-value cutoff was applied. For conservation analysis, the phastCons 60-way bigwig track for mm10 was downloaded from UCSC genome browser (http://genome.ucsc.edu, (Nassar et al., 2023)). Peaks were stratified based on how many datasets they were present in. Peaks in each group were treated as bins and the average conservation score within each peak-bin was calculated using deepTools computeMatrix, and then the output matrix was used for plotting and the values for statistical analysis, which was done using a One-way ANOVA followed by Tukey’s post-hoc adjustment. UCSC genome browser was also used for visualizations in Figure 5G.

### Comparison with Published Datasets

Peak regions and if applicable bigwig files for ChIP-seq and CUT&RUN datasets were downloaded from the references given. If necessary, peak lists were converted from mm9 to mm10 using the UCSC LiftOver (Hinrichs et al., 2006) with default settings. RNA-seq bulk RPKM data was downloaded from the reference, and these values were plotted for each target. The preprocessed single-cell RNA sequencing (scRNAseq) dataset from the mouse embryonic limb atlas (10x_limbdataset) (He et al., 2020), was obtained from the UCSC single-cell browser (https://mouse-limb.cells.ucsc.edu/) as a scanpy-compatible 10x.h5ad file. The Anndata objects were analyzed using Python packages Scanpy (v.1.9.5) (Wolf et al., 2018) and Anndata (v.0.9.2) (https://github.com/scverse/anndata), inspired by Seurat (Butler et al., 2018). The dataset was filtered for stage E.11, containing 17,574 cells, and the gene expression of interest was visualized using UMAP coordinates matching the original paper. Cell type cluster assignments were based on cell labels and marker genes from the original study.

## Data Availability

Raw data, normalized bigwig files, and peak lists have been uploaded to ArrayExpress, and are awaiting accession number E-MTAB-14340. Published datasets used for comparison can be downloaded according from the references listed.

## Supplementary Figures and Tables

**Figure S1.**
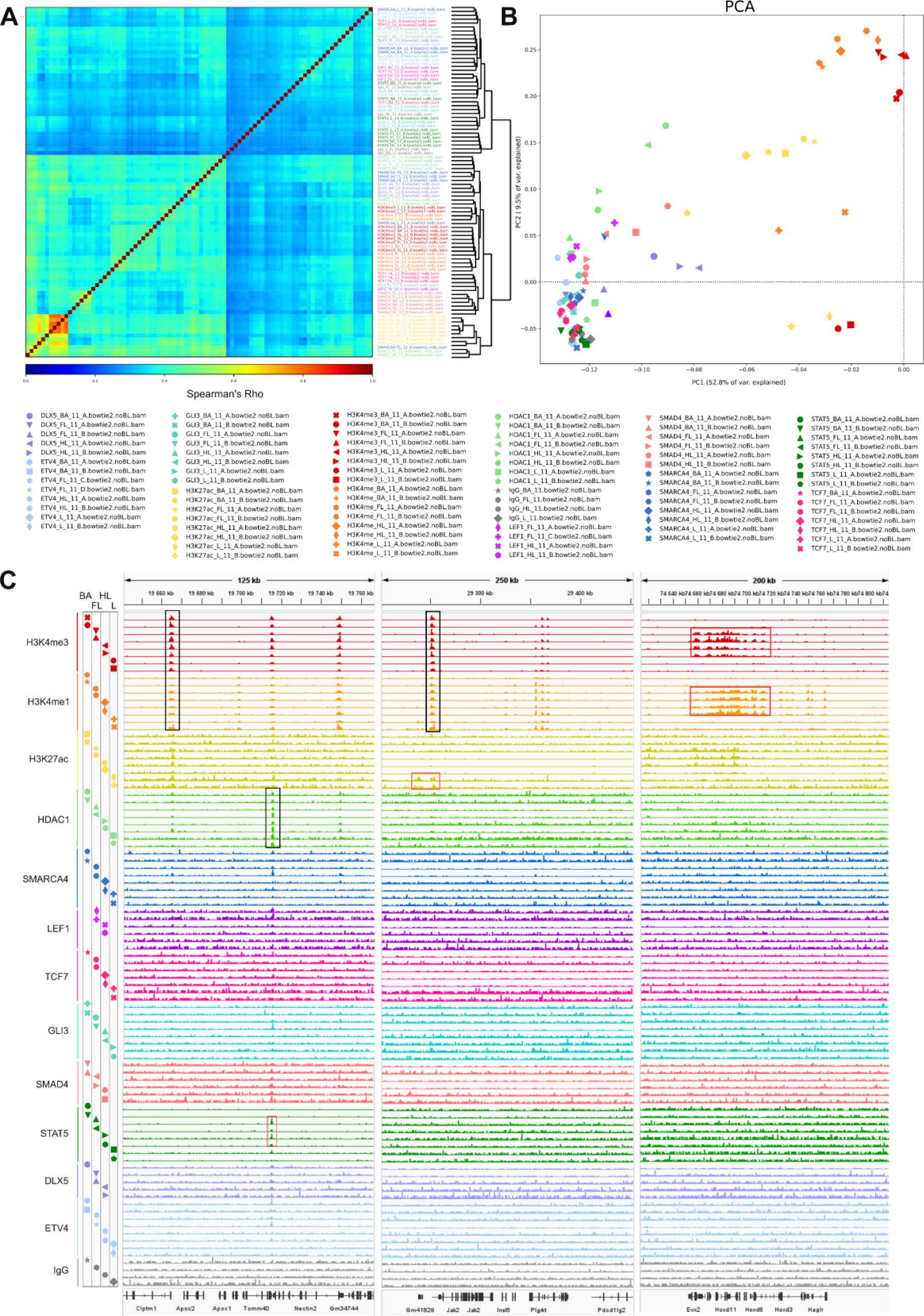
A. Spearman’s correlation plot of all C&R replicate datasets. Sample names are colored by target. **B.** PCA analysis of all C&R replicate datasets, data points are colored by target and full dataset names are included in the legend below. **C.** C&R datasets visualized in IGV, showing loci Apoc1, Jak2 and Hoxd. Common peaks by target are shown with black boxes, tissue specific instances are shown in red. Replicates are normalized to reads per genome coverage and are group scaled by factor. BA = branchial arches, FL = forelimb, HL = hindlimb, L = liver.

**Figure S2.**
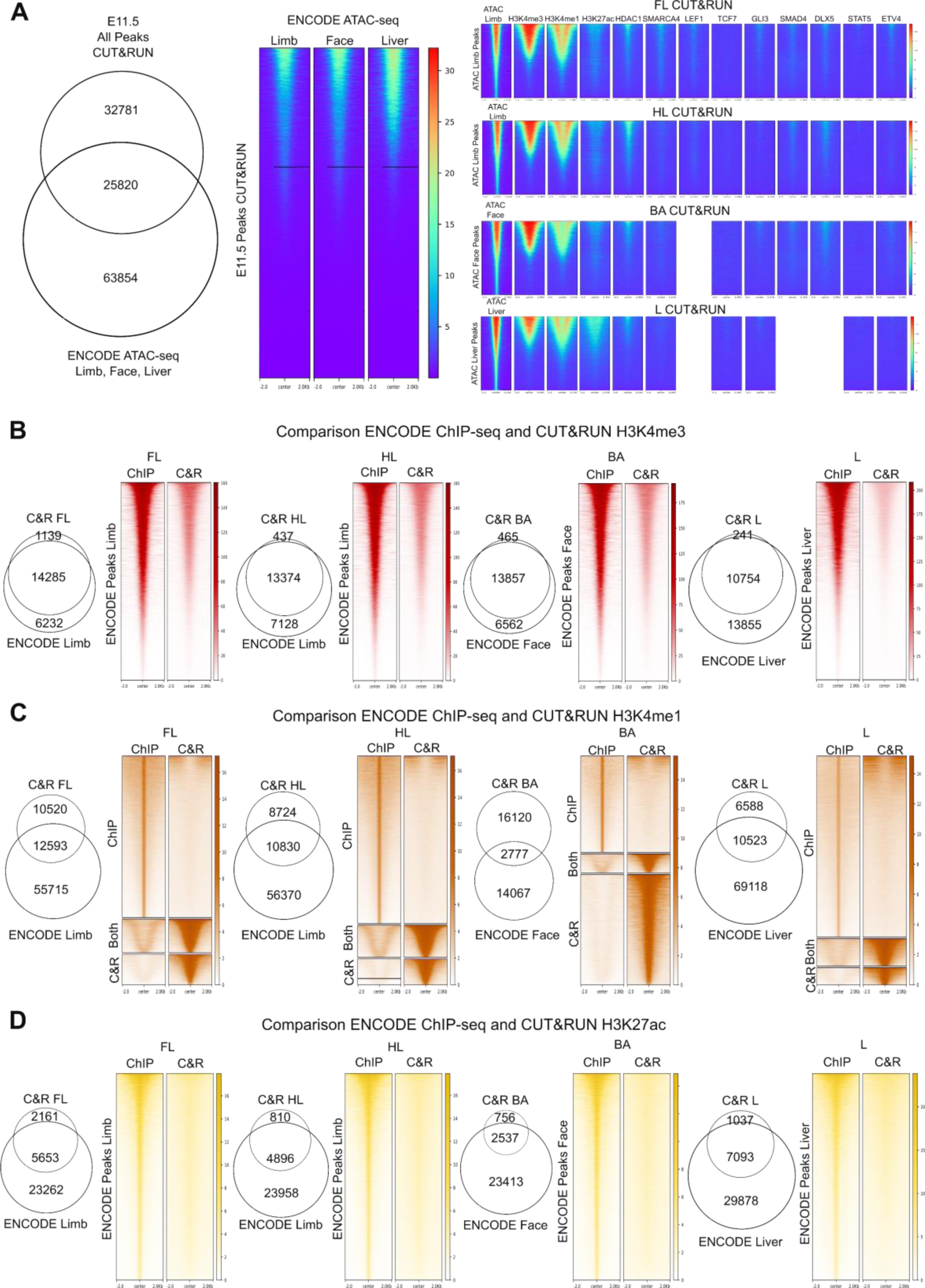
A. Left: Venn diagram comparing all E11.5 C&R peaks to open chromatin ENCODE ATAC-seq in the limb, liver and facial prominence (face) at E11.5. Middle: heatmaps showing ATAC-seq signal within C&R peaks. Right: Heatmaps showing C&R signal (top to bottom: FL HL BA L) within ATAC-seq peaks. B. Comparison of ENCODE H3K4me3 data in the limb, face, and liver to C&R data including venn diagrams and signal plots, showing high similarity between the C&R and ChIP datasets. C. Comparison of ENCODE H3K4me1 data to C&R. H3K4me1 showed a much lower overlap with ChIP: the overlapping peaks between the C&R and ChIP had a broader signal profile, and the unique C&R peaks had a similar shape. In contrast, the ChIP datasets contained many very sharp peaks which were almost completely devoid of signal in our C&R. D. Comparison of ENCODE H3K27ac data to C&R showing a high similarity between the two datasets, with more peaks being called in the ChIP, but many of these regions containing signal also in the C&R.

**Figure S3.**
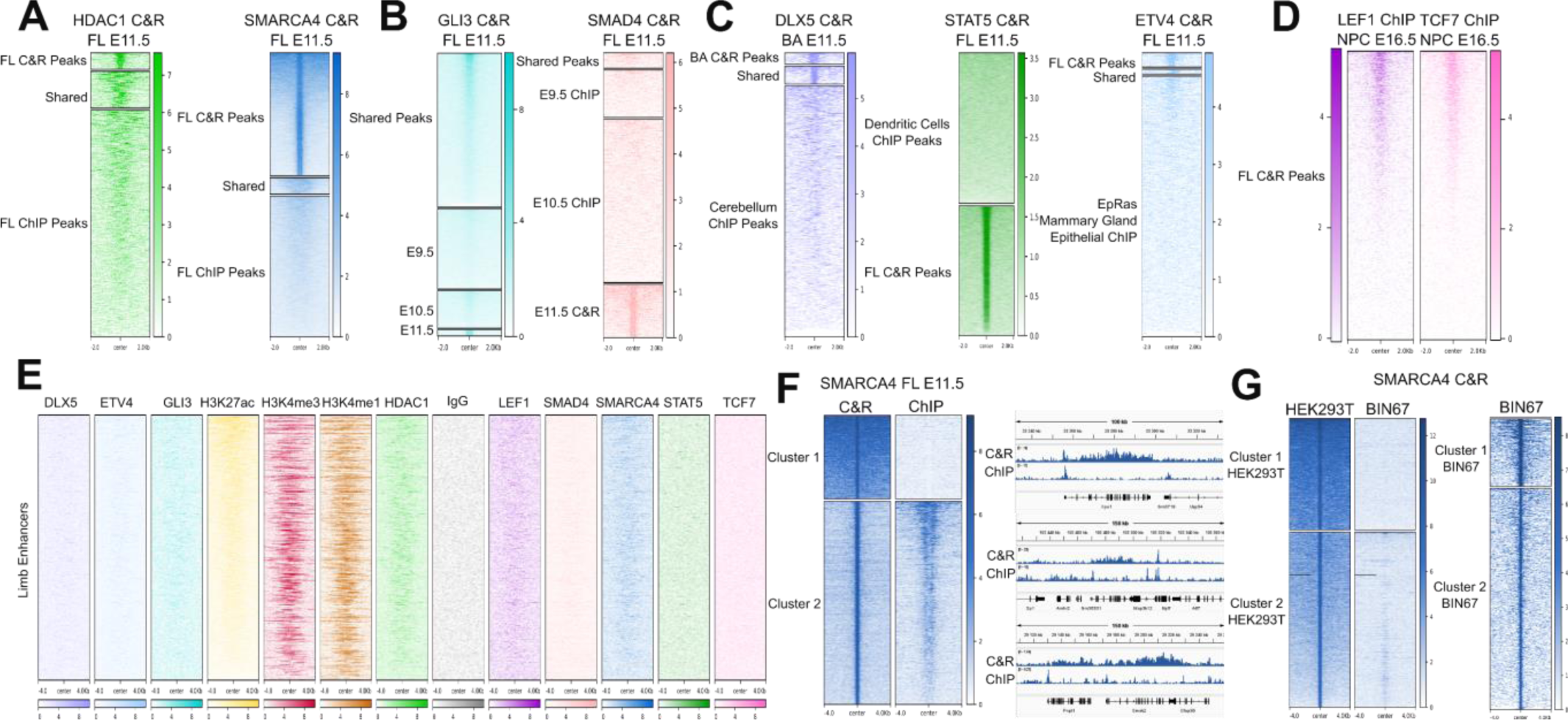
A. Heatmaps of HDAC1 (left) and SMARCA4 (right) within peaks defined by C&R, those shared with matching ChIP-seq datasets, and ChIP specific peaks. B. Heatmaps of GLI3 (left) and SMAD4 (right) C&R signal within peaks defined by published datasets at E9.5 and 10.5 (GLI3 C&R, left and SMAD4 ChIP-seq, right) and our E11.5 peaks, and those shared between timepoints. GLI3 peaks and signal were more consistent over time compared to SMAD4 (Figure S3B). Interestingly the GLI3 datasets were also C&R, versus the SMAD4 datasets were performed with ChIP-seq, so this might reflect a technological instead of biological observation. C. Left: heatmaps showing DLX5 C&R signal within BA specific peaks, cerebellum ChIP-seq peaks, and shared. Center: heatmaps showing STAT C&R signal in dendritic cell ChIP-seq peaks and in FL C&R peaks. Overlapping peaks (28) were too little to be shown. Right: heatmaps of ETV4 C&R signal within FL C&R peaks, EpRas ChIP-seq peaks, and shared peaks. These datasets, being less comparable to our C&R (as far as tissue and timepoint), logically showed less of a signal overlap. D. Heatmaps showing ChIP-seq signal for LEF1 and TCF7 in nephron progenitor cells, within peaks called for LEF1 and TCF7 in FL C&R. Since peak sets weren’t provided, we instead look at signal from the NPCs in our FL peaks. Interestingly, we found that nearly half of FL peaks had some signal enrichment in the NPCs, indicating some conservation of Wnt target genes between the developing limb and kidney. E. Heatmaps showing signal profiles for all C&R targets within the limb enhancers defined by Andrey et al.. The majority of these enhancers were marked by H3K4me3, H3K4me1, and H3K27ac, while we found less overall enrichment in our ChR and TFs. F. Left: Heatmaps showing C&R and ChIP-seq signal targeting SMARCA4 in E11.5 FL within clusters 1 and 2 of C&R FL peaks. Right: IGV visualization of C&R versus ChIP signal profiles. Notably, these broadly enriched regions were not enriched in ChIP-seq of SMARCA4 in the same tissue, though some of the narrow peaks are similar between the datasets. G. Heatmaps showing SMARCA4 C&R signal profiles from HEK293T cells and published data in BIN27 cells, within clustered peaks called from the HEK293T data (left) and BIN67 signal within its own peaks (right). To explore if broad peaks was a C&R specific phenomenon, or possibly an antibody specific one, we looked at a published C&R done in human ovarian small cell carcinoma cells (BIN67) with SMARCA4 overexpression (Orlando et al., 2020) and performed a SMARCA4 C&R in the human cell line HEK293T. We observed the same phenomenon of broad peaks in our C&R, though this was not replicated in the published data (Figure S3G), indicating that it could be related to the antibody used, though we cannot rule out cell-type or protocol specific causes.

**Figure S4.**
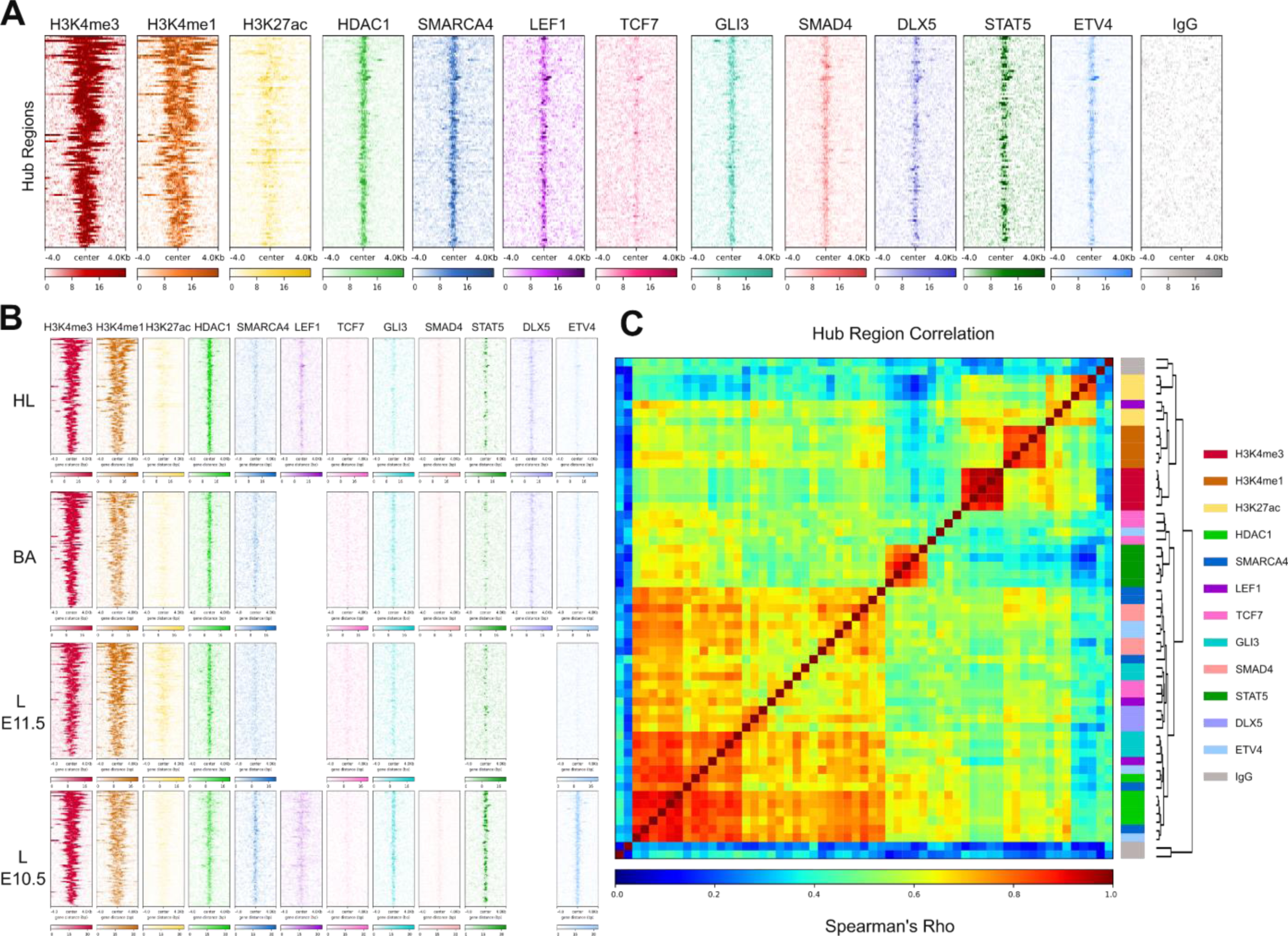
A. Heatmaps showing signal intensity for each target within the 99 hub regions. B. Heatmaps showing signal intensity in hub regions for the targets in the other tissues and time points. C. Spearman’s correlation of all datasets within the hub regions, showing that even within the highly bound regions, target specific idiosyncrasies lead to many instances of higher correlations based on target. Dendrogram connects to boxes that are color coded based on target, see for examples STAT5 (dark green) and HDAC1 (light green).

**Table S1.** Antibodies Information.

| Name | Code | Batch/Lot |
| --- | --- | --- |
| H3K4me3 | ABIN6971977 | 30720008 |
| H3K4me1 | ABIN3023251 | 3560036504 |
| H3K27ac | ABIN6971893 | 06921014 |
| HDAC1 | ABIN2854776 | NA |
| SMARCA4 | ABIN6991990 | 7749-1303 |
| TCF7 | ABIN5620945 | X22061404 |
| LEF1 | ABIN1680678 | 5500007091 |
| GLI3 | ABIN2855813 | 40716 |
| SMAD4 | ABIN6972727 | 34614001 |
| DLX5 | ABIN5619487 | X21080310 |
| STAT5 | ABIN5557532 | BA7271967 |
| ETV4 | ABIN6131217 | 74340301 |

